# Gain and loss of plasmid-borne antibiotic resistance genes are associated with chromosomal resistance presence in Enterobacteriaceae

**DOI:** 10.1101/2025.10.17.683045

**Authors:** Yang Liu, Yue Liu

## Abstract

Plasmids are central vehicles for the dissemination of antibiotic resistance genes (ARGs). They are among the most mobile and evolvable genetic elements, with broad host ranges and high rates of gene turnover, making them especially effective in spreading antibiotic resistance across bacterial lineages. Using the phylogeny-aware gene gain and loss model applied to 6,895 Enterobacteriaceae genomes, we quantified four evolutionary processes—gene gain, loss, expansion, and reduction—for plasmid-borne genes. We found that, overall, plasmid-borne ARGs (pARGs) exhibit similar gain rates compared with other plasmid genes, but significantly higher expansion and reduction rates. All four processes were strongly species dependent, with only a minor influence of antibiotic class. Further, bacterial clades harboring chromosomal ARGs (cARGs) showed significantly higher acquisition and lower loss of plasmid-borne resistance than did their sister clades lacking corresponding cARGs. Moreover, we found that the IncQ2 backbone was associated with *qnrS2* and exclusively identified in *Leclercia adecarboxylata*, while Col(VCM04) plasmids carrying *mprF* were predominantly (71.4%) distributed within the *Citrobacter* genus. In summary, plasmid-mediated resistance is primarily species-dependent, and cARGs effectively mark lineages with a high capacity for plasmid-borne resistance acquisition.

**Importance:** Plasmids play a central role in the spread of antibiotic resistance genes (ARGs), and the long-term evolutionary behavior of plasmid-borne ARGs (pARGs) could provide insights into the emergence of novel multidrug resistance. We studied nearly 7,000 Enterobacteriaceae genomes and show that pARGs evolve through the same gain processes as other plasmid genes but exhibit markedly higher and species-dependent copy number changes. Crucially, the strong association between chromosomal and plasmid ARGs reflect a lineage-level pattern of resistance retention, likely shaped by historical selective pressures or specific genomic backgrounds. Identifying such evolutionarily lineages may provide a basis for predicting and monitoring the emergence of multidrug resistance.

## INTRODUCTION

Antibiotic resistance is a major, ongoing challenge in both medicine and evolutionary biology (1). In bacterial genome evolution, gene gain through horizontal gene transfer (HGT) and gene loss (via selection) are the primary forces shaping genomic repertoires (2, 3). Mobile genetic elements (MGEs), particularly conjugative plasmids, serve as principal vehicles of HGT and are central to the spread of antibiotic resistance genes (ARGs) (4–6).

Comparative genomic analyses show that plasmid taxonomic units carrying ARGs exhibit significantly higher mobility, broader host ranges, and elevated rates of homologous recombination and gene turnover, indicating that resistance dissemination is facilitated by intrinsically plastic and mobile plasmid lineages (7, 8). Their plasticity is enhanced by high densities of transposable elements and recombination systems, which accelerate gene repertoire turnover (9, 10). Moreover, evolutionary innovations such as replication-control mutations can increase plasmid copy number, thereby amplifying resistance levels and enhancing transfer efficiency (11). Thus, plasmids are highly dynamic replicons characterized by frequent horizontal transfer, rapid turnover, and extensive genetic plasticity (12, 13).

Given their evolutionary plasticity, understanding plasmid-mediated ARG turnover is particularly critical in clinically important bacterial families such as Enterobacteriaceae (12). The Enterobacteriaceae family harbors extensive plasmid diversity and constitutes one of the largest reservoirs of clinically relevant ARGs (14–16). Members of this family, including *Escherichia*, *Klebsiella*, *Enterobacter*, and *Salmonella*, are major contributors to multidrug-resistant infections in hospitals and communities worldwide (17). Although numerous clinical studies have documented outbreaks of antibiotic resistance within the Enterobacteriaceae family (16–18), most of these reports have focused on the dissemination of particular resistance plasmids or the epidemiology of individual ARGs (19, 20). However, key questions remain as to how plasmid-borne ARGs in Enterobacteriaceae are maintained and amplified across lineages, and whether their evolutionary dynamics differ from those of other plasmid-encoded functions.

Here, we systematically investigate ARG turnover dynamics in Enterobacteriaceae plasmids by applying a phylogeny-aware birth-death model to 6,895 genomes. Specifically, we (i) quantify the balance of ARG gain, loss, expansion, and reduction across diverse species, (ii) assess the association between chromosomal and plasmid-borne resistance repertoires.

## RESULTS

### Plasmid-borne antibiotic resistance genes show higher expansion and reduction rates but similar gain dynamics compared with other plasmid genes

To investigate the evolutionary dynamics driving the current distribution of antibiotic resistance genes (ARGs) in Enterobacteriaceae plasmids, we applied a phylogenetic birth-death model using Count (21) to quantify gene turnover rates across 6,895 genomes of which 5,335 contained at least one identified plasmid sequence (Supplementary Table S1). The approach estimates four key evolutionary processes: (ⅰ)gene family turnover, comprising gain (acquisition of new gene families) and loss (complete disappearance of an existing family) (Supplementary Figure S1A, C); and (ⅱ)copy-number variation, comprising expansion (increase in gene copies within an existing family) and reduction (decrease in gene copies while maintaining at least one copy) (Supplementary Figure S1B-C). Since we focused exclusively on plasmid-associated gene families (defined by a ≥ 70% plasmid-occurrence threshold), expansion can be biologically interpreted as intra- or inter-plasmid gene duplication (e.g., mediated by transposons), the acquisition of additional plasmid carrying the same gene family, or even chromosomal integration of a plasmid-borne gene copy, while reduction represents the subsequent loss of these redundant copies (Supplementary Figure S1B).

We compared overall rates between plasmid borne ARGs (pARGs) and all other plasmid genes (non-ARGs) across 80 species. Our analysis revealed that pARGs exhibit substantially different dynamics for gene gain and loss versus expansion and reduction (Figure 1, Supplementary Table S2). Gain rates showed no significant differences between pARGs and non-ARGs (n = 80, gain: adjusted p = 0.257; Figure 1A), with data points forming relatively compact distributions across species. This suggests that the fundamental processes governing the acquisition of entire gene families operate similarly for resistance and non-resistance genes. In contrast, pARGs exhibited significantly elevated rates for loss (n = 80, adjusted p = 1.23×10^−2^, median log₂(ARG/ALL) = 0.15; Figure 1B), expansion (adjusted p = 3.34×10^−6^, median log₂(ARG/ALL) = 1.28; Figure 1C) and reduction (adjusted p = 4.03×10^−5^, median log₂(ARG/ALL) = 0.83; Figure 1D) events. The high scatter in these data points indicates substantial species-specific variation in the extent to which loss, expansion and reduction dynamics differ between pARGs and other plasmid genes. Regarding the ratios of gain-to-loss and expansion-to-reduction, no significant differences were observed for pARGs (p > 0.05; Supplementary Figure S2).

**Figure 1.**
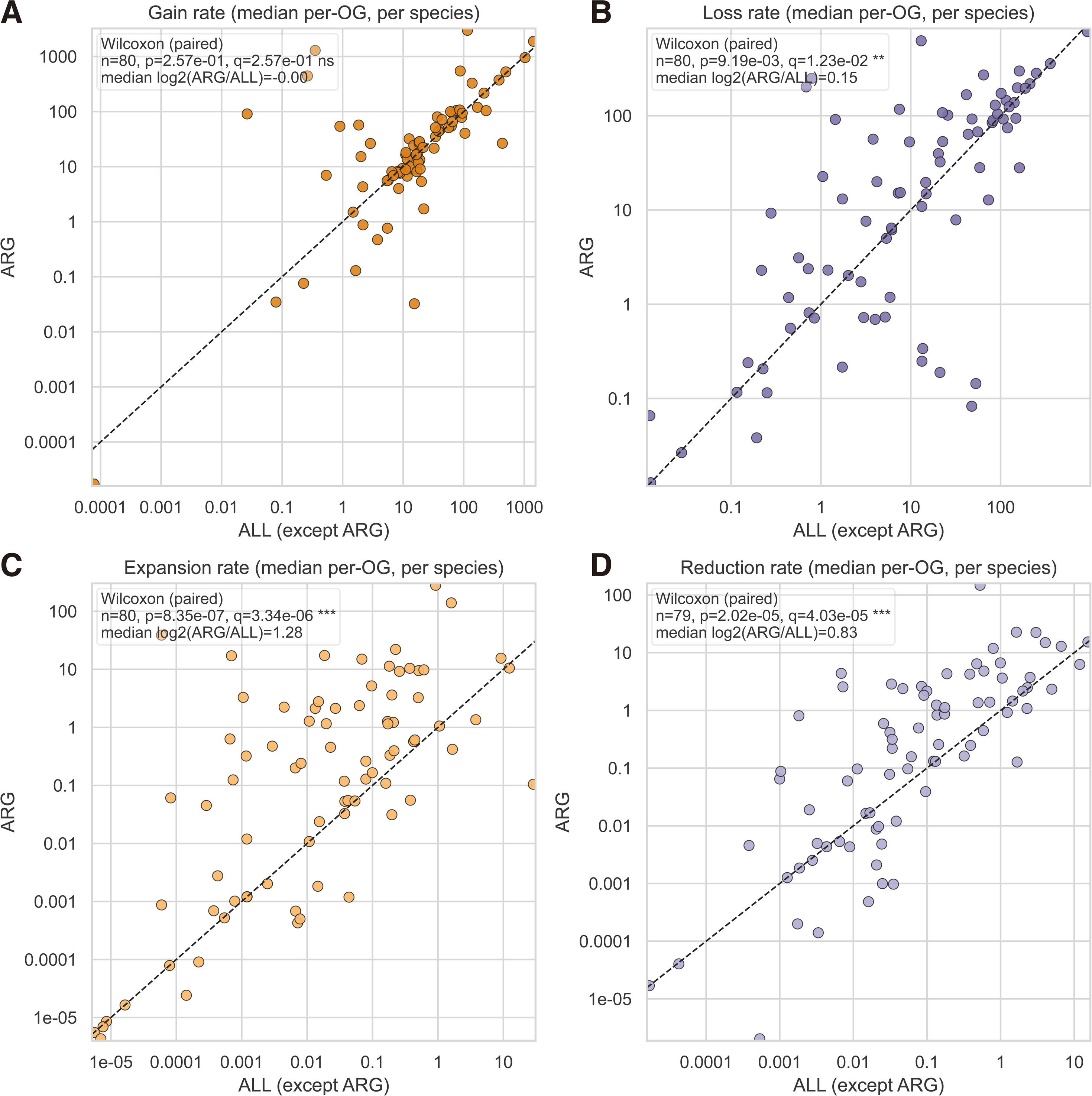
Evolutionary dynamics of plasmid-borne antibiotic resistance genes compared to other plasmid genes in Enterobacteriaceae. (A) Gain rates (appearance of new gene families per substitution per site) for ARGs versus other plasmid genes. (B) Loss rates (disappearance of gene families per substitution per site) for ARGs versus other plasmid genes. (C) Expansion rates (addition of gene copies within existing families per substitution per site) for ARGs versus other plasmid genes. (D) Reduction rates (loss of gene copies within existing families per substitution per site) for ARGs versus other plasmid genes. Gene families were defined as orthologous groups (OGs).

In summary, across Enterobacteriaceae, gain rates show no pARG-specific signal, indicating that acquisition of pARG families follows common plasmid evolutionary pathways. In contrast, pARGs exhibit significantly elevated expansion and reduction rates with pronounced species-level variation, revealing that these copy-number dynamics are primarily driven by species identity. While gene loss rates showed a statistically significant difference, the effect size was small (median log2 ratio = 0.15).

### Turnover dynamics of antibiotic resistance genes are primarily species dependent

To test whether the species-level patterns in Figure 1 are consistent across resistance mechanisms or vary among drug classes, which are defined by their mechanisms of action, we assigned orthologous gene families to CARD-defined drug classes and estimated gain, loss, expansion, and reduction rates for each class across species. Using the CARD database (22), we identified and analyzed the turnover dynamics of specific drug classes, which are groups of antibiotics with similar chemical structures and mechanisms of action.

Our analysis revealed that the evolutionary rates for all four event types were strongly species-dependent (Kruskal–Wallis test; gain: H = 536.9, p = 2.9×10^−69^, η² = 0.76; loss: H = 506.3, p = 1.4×10^−63^, η² = 0.71; expansion: H = 312.5, p = 1.6×10^−29^, η² = 0.39; reduction: H = 436.9, p = 2.4×10^−51^, η² = 0.60), with substantial variation across species for gain, loss, expansion, and reduction (Supplementary Figure S2A–B; Figure 2A–B). In contrast, drug class effects were weak: no significant signals were detected for gain (p = 0.74) and loss (p = 0.58), while expansion (p = 5.2×10^−9^, η^2^=0.09) and reduction (p = 1.1×10^−3^, η^2^=0.04) showed significant but weak drug class effects, which were an order of magnitude smaller than those of species. To account for potential biases from multi-counting genes across drug classes, we used an Ordinary Least Squares (OLS) model for variance partitioning. This analysis confirmed species identity as the primary driver of pARG turnover, explaining 66.8%, 63.1%, 42.1%, and 51.5% of the total variance for gain, loss, expansion, and reduction events, respectively. In contrast, drug class identity accounted for minimal variance across all events: 2.1% for gain, 1.7% for loss, 6.9% for expansion, and 4.1% for reduction. Full results are provided in Supplementary Table S3.

**Figure 2.**
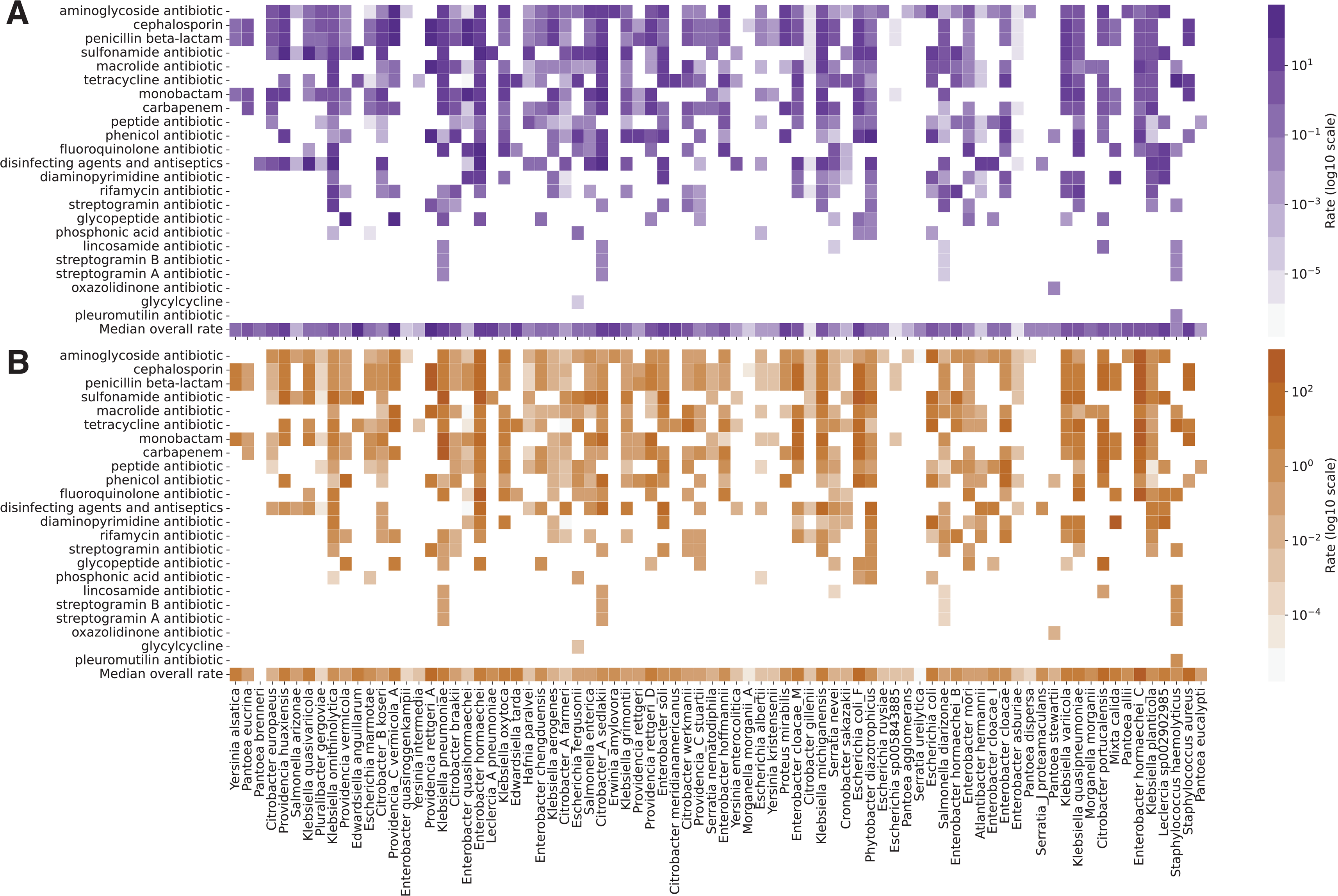
Expansion and reduction rate patterns across antibiotic resistance classes and Enterobacteriaceae species. Heatmaps showing evolutionary rates for different antibiotic resistance classes across bacterial species. (A) Expansion rates. (B) Reduction rates. Rows represent Enterobacteriaceae species, and columns represent CARD-defined antibiotic resistance classes. Color intensity indicates log10 scaled rate magnitude, with darker colors representing higher rates. White cells indicate absence of the ARG class in the species.

Furthermore, species that exhibited high expansion rates typically also showed elevated reduction rates for the same drug classes (n=80, Spearman’s rank correlation with 5,000 permutations: ρ=0.792, permutation p≈0.0002). By contrast, species with high gain rates did not necessarily display the corresponding high loss rates (n=80, ρ=-0.196, p≈ 0.079), indicating that plasmid-borne gene expansion and reduction increases and decreases within existing gene families are more tightly coupled than complete gene family gain and loss.

In summary, antibiotic resistance gene evolution dynamics are primarily driven by species identity, with only minor drug class effects. Moreover, expansion and reduction rates are coupled, whereas gain and loss are largely independent.

### Chromosomal antibiotic resistance genes (cARGs) are associated with elevated pARG acquisition rates

Our previous analyses revealed that pARGs exhibit similar gain rates compared to other plasmid genes. We further investigated whether additional factors might be associated with the evolutionary dynamics of pARGs, specifically examining whether the presence of antibiotic resistance genes in bacterial chromosomes is associated with altered evolutionary rates of pARGs. We employed a comparative phylogenetic approach using sister clade analysis (Figure 3A), identifying 464 pairs of phylogenetic clades where one clade (lineage) contained ≥80% of genomes harboring a specific ARG family in their chromosome (ARG-containing clade) while <20% of genomes in the sister clade (lineage) contained that homologous ARG family. For each clade pair, we calculated gain and loss rates for plasmid genes by summing evolutionary events across all branches within each clade and normalizing by total branch length. In total, 80 species and 84 RGI-defined ARG families were included in the analysis (Supplementary Tables S4 and S5). Linear mixed-effects models (LMM) were used as a statistical test for the analysis, including fixed effects for clade type, gene class, and their interaction, with random effects for species, ARG family, and sister-clade comparisons to capture lineage-specific variation (see Methods).

**Figure 3.**
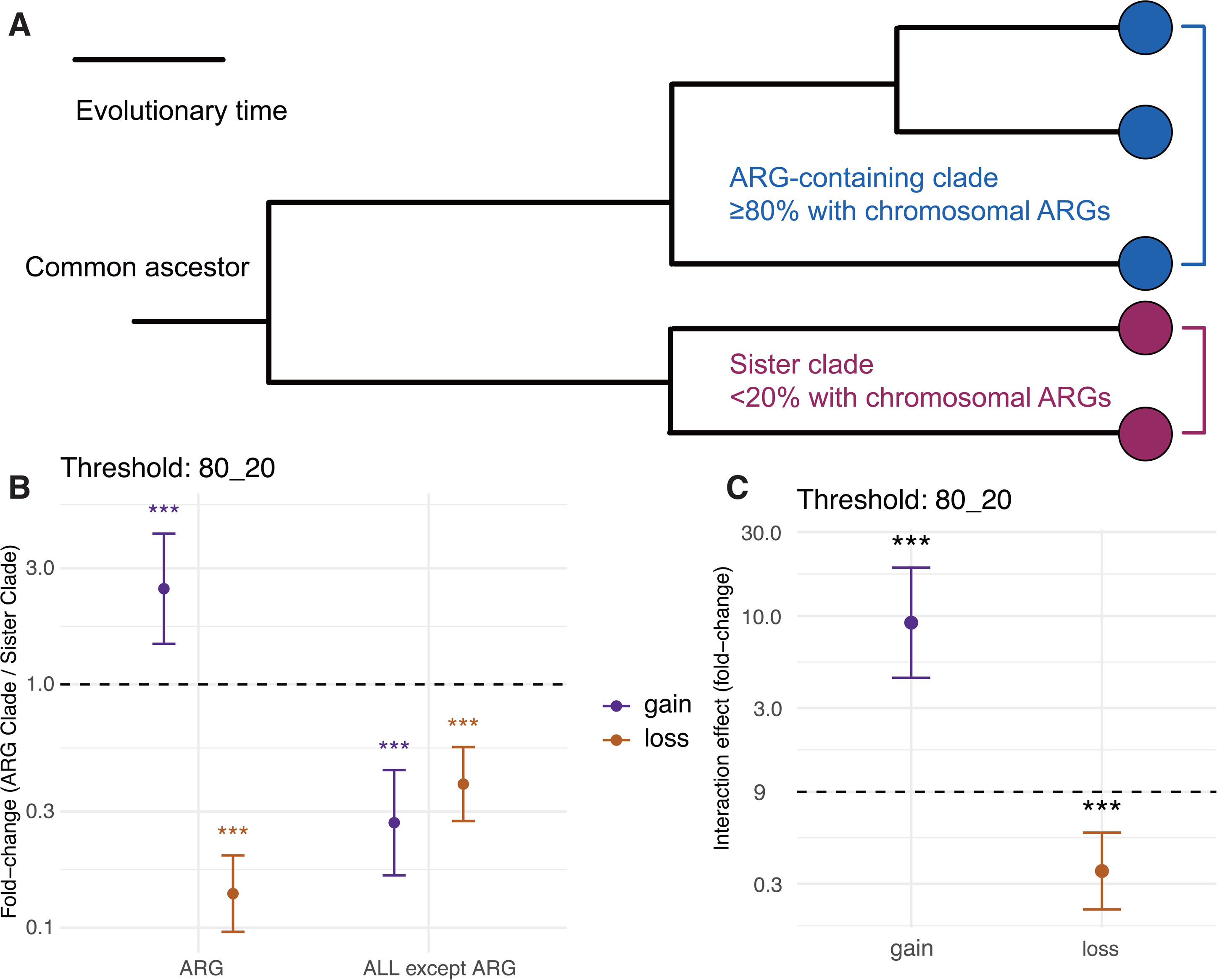
Association between presence of cARGs in bacterial lineages and plasmid borne ARG acquisition. (A) Schematic showing sister clade comparison approach. Phylogenetic clades containing high frequencies (≥80%) of specific cARG families are compared to sister clades with low frequencies (<20%). Colored circles represent different bacterial clades. (B) Fold-change comparison of evolutionary rates between ARG-containing clades and sister clades. Y-axis shows the fold-change ratio (ARG clade / Sister clade) for gain (purple) and loss (orange) rates of plasmid ARGs versus other plasmid genes. Dashed line at y=1 indicates no difference between clades. (C) Interaction effect showing the differential impact of cARG presence on gain versus loss dynamics. Y-axis represents the interaction effect (fold-change, log scale) between clade type and evolutionary process. Error bars show 95% confidence intervals. Statistical significance indicators: ***p < 0.001, *p < 0.05, ns = not significant.

Our analysis revealed that bacterial clades with cARGs exhibit significantly different plasmid gene gain rates compared with their sister clades (Figure 3B-C). ARG-containing clades showed approximately 2.47-fold higher gain rates for pARGs (estimate = 0.91 on the log scale, p = 1.9×10⁻⁷; Figure 3B, Supplementary Table S6) and significantly lower loss rates (fold-change = 0.138, p = 0.034). In contrast, other plasmid genes showed significantly lower gain rates in ARG-containing clades compared with sister clades (about 3.7-fold lower; estimate = −1.3, p = 2.97×10^−7^), while their loss rates also showed a decrease between clade types (estimate = −0.94, p = 1.44×10^−7^). The interaction term indicated that the correlation between cARG presence and evolutionary rates differs significantly between gene types for both gain (estimate = −2.2, p = 2.2 × 10^−9^) and loss events (estimate = −1.0, p = 5.6 × 10^−5^) (Figure 3C, Supplementary Table S6).

To test the robustness of our results across different clade definitions, we performed an analysis across a gradient of sister-clade thresholds (70/30, 80/20, 85/15, and 90/10 for cARG/sister clades). Although the magnitude of the effect was sensitive to the specific cutoff, the positive association remained robust across all thresholds (Supplementary Figure S4, Supplementary Table S5). As the threshold for defining ARG-containing clades became more stringent, representing lineages with higher levels of chromosomal resistance prevalence, the magnitude of the pARG gain bias became more pronounced. Specifically, the fold-change in pARG gain rates increased progressively from 1.65 at the 70/30 threshold (p = 0.015) to 2.47 at the 80/20 threshold (p = 6.7 × 10^−4^), 3.21 at the 85/15 threshold (p = 1.5 × 10^−4^), and 4.91 at the 90/10 threshold (p = 1.19 × 10^−6^) (Supplementary Figure S4, Supplementary Table S6). Parallel to the gain dynamics, the suppression of pARG loss rates in cARG clades remained remarkably stable and highly significant across all tested categories. The loss rate fold-changes ranged from 0.183 to 0.125 with all *p* values below 0.001 (Supplementary Figure S4, Supplementary Table S6), indicating that these lineages consistently maintain acquired resistance genes more effectively than their sister clades.

In addition to the overall association, we performed per-family LMM analyses for those with at least three available clades in Enterobacteriaceae (Supplementary Table S7). The overall gene gain rate remained higher in cARG clades (Supplementary Figure S5; p < 0.05). Specific ARG families, such as sulfonamide-resistant *sul* and trimethoprim-resistant dihydrofolate reductase (*dfr*), display higher gene gain but lower gene loss rates (Supplementary Figure S6).

Notably, individual tests for each family revealed that a larger proportion of cARG families exhibited significantly reduced gene loss rates than elevated gain rates (see Supplementary Figure S6). Along with the reduced loss of other plasmid genes within these lineages (Supplementary Figure S3), this stability may represent an evolutionary tendency to maintain plasmids in cARG-containing clades. Consistent with this, a pairwise comparison of 436 clade pairs revealed that cARG clades maintain a significantly higher plasmid burden than their sister counterparts, with a median of 2.64 unique plasmid clusters compared to 1.66 (*p* = 2.42 × 10^−21^) (Supplementary Figure S7A).

To further study whether the accelerated pARG acquisition in cARG clades be attributed to lower plasmid backbone incompatibility, we then calculated the frequency of these pARG-associated replicon types within their respective sister clades. We observed a 0.00% median sharing ratio of backbone replicon genes between sister clades (Supplementary Figure S7B), suggesting that most of these pARG-carrying plasmids were absent in the most recent common ancestor and were acquired specifically in the cARG clades. Together, these evidences help strengthen the interpretation that these cARG-containing lineages are genuinely more prone to plasmid-mediated resistance acquisition.

For most species, cARG-clade associated genomes exhibit a wide and diverse spectrum of antibiotic resistance (Figure 4A). Many species (e.g., *Enterobacter cloacae_M*, *Enterobacter chengduensis*, *Erwinia persicina*, *Klebsiella terrigena*, and *Kosakonia cowanii*) exhibit widespread resistance to clinically and agriculturally used antibiotics (e.g., penicillins, cephalosporins, and tetracyclines). In contrast, only a few species accumulate antibiotic-resistant strains for a specific antibiotic; the most notable examples are *Serratia nevei* and *Klebsiella michiganensis*, which exhibit high levels of aminoglycoside antibiotic resistance, and *Klebsiella quasipneumoniae* for glycopeptide antibiotics. The observed resistance profiles across these lineages can be compared to the representative consumption levels of corresponding antibiotic classes in clinical and agricultural settings (Figure 4B and 4C).

**Figure 4.**
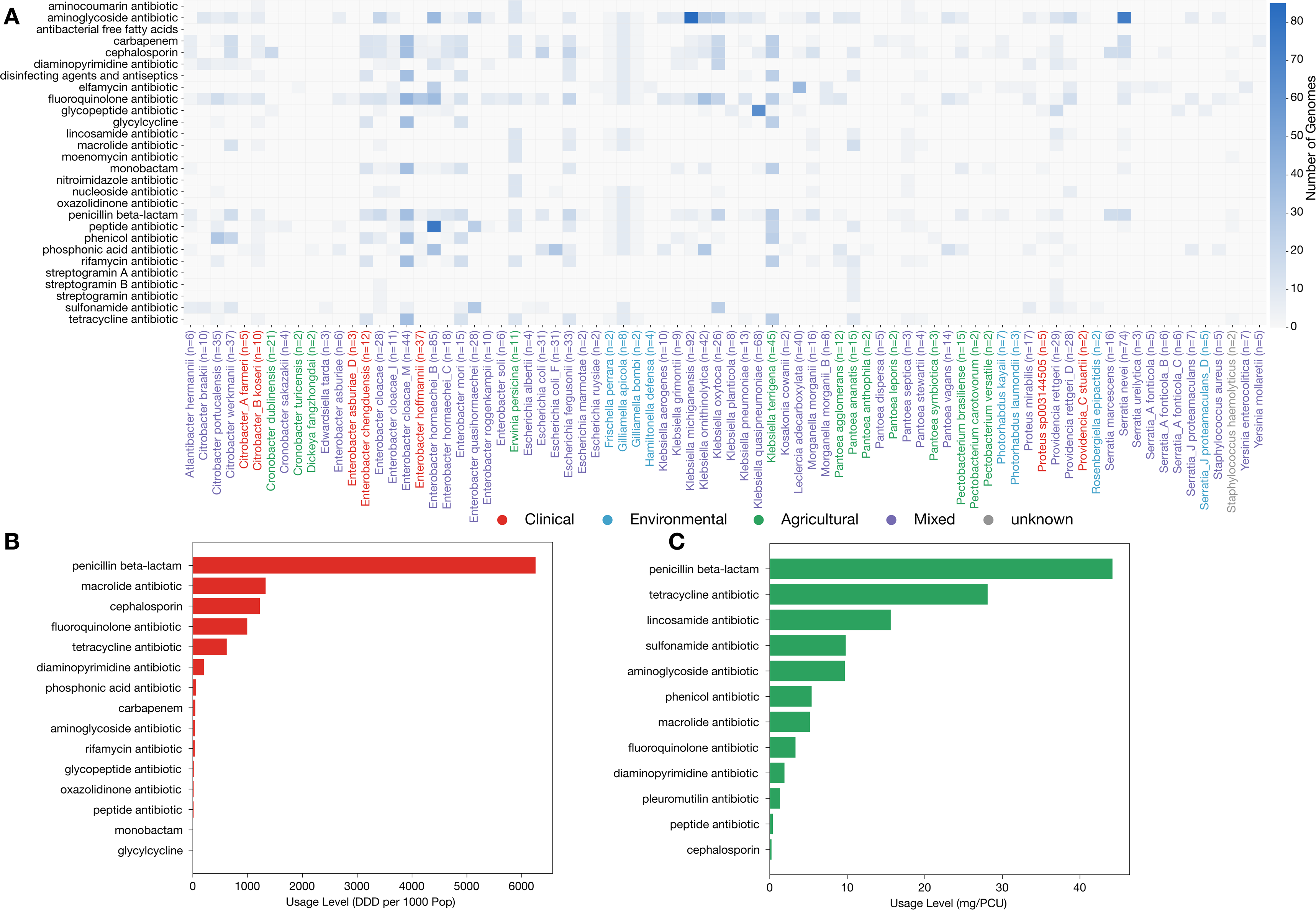
ARG distribution within cARG-associated lineages and antibiotic consumption levels. (A) Heatmap of ARG profiles across cARG-containing clades. The x-axis labels denote the bacterial species and total genome numbers within each analyzed clade. Species names are color-coded according to their primary habitat: Clinical (red), Environmental (blue), Agricultural (green), Mixed (purple), or Unknown (grey). The y-axis lists antibiotic drug classes as defined by the Comprehensive Antibiotic Resistance Database (CARD). Color intensity indicates the number of genomes harboring specific resistance gene classes. (B) Clinical antibiotic consumption. Bar chart representing human antibiotic usage data from Spain (2023). The x-axis measures consumption in Defined Daily Doses (DDD) per 1,000 people per day. (C) Agricultural antibiotic consumption. Bar chart representing veterinary antimicrobial sales data from Spain (2022). The x-axis measures sales in milligrams per population correction unit (mg/PCU). Antibiotic categories are mapped into standardized CARD drug classes for cross-comparison with the heatmap.

### Co-occurrence of plasmid-borne antibiotic resistance genes with plasmid replicon types

To identify the associations between the presence of ARGs and specific plasmid types, we reconstructed plasmid units from draft assemblies and performed replicon typing for both draft and complete assemblies. For plasmid groups defined by a particular replicon type, we calculated all-against-all Mash distances and constructed plasmid similarity networks. Notably, we employed a Mash distance cutoff of 0.025, which is equivalent to the secondary clustering criterion used by MOB-recon tool. Consequently, each panel in Figure 5 visualizes the fine-grained genetic similarity within specific plasmid types.

**Figure 5.**
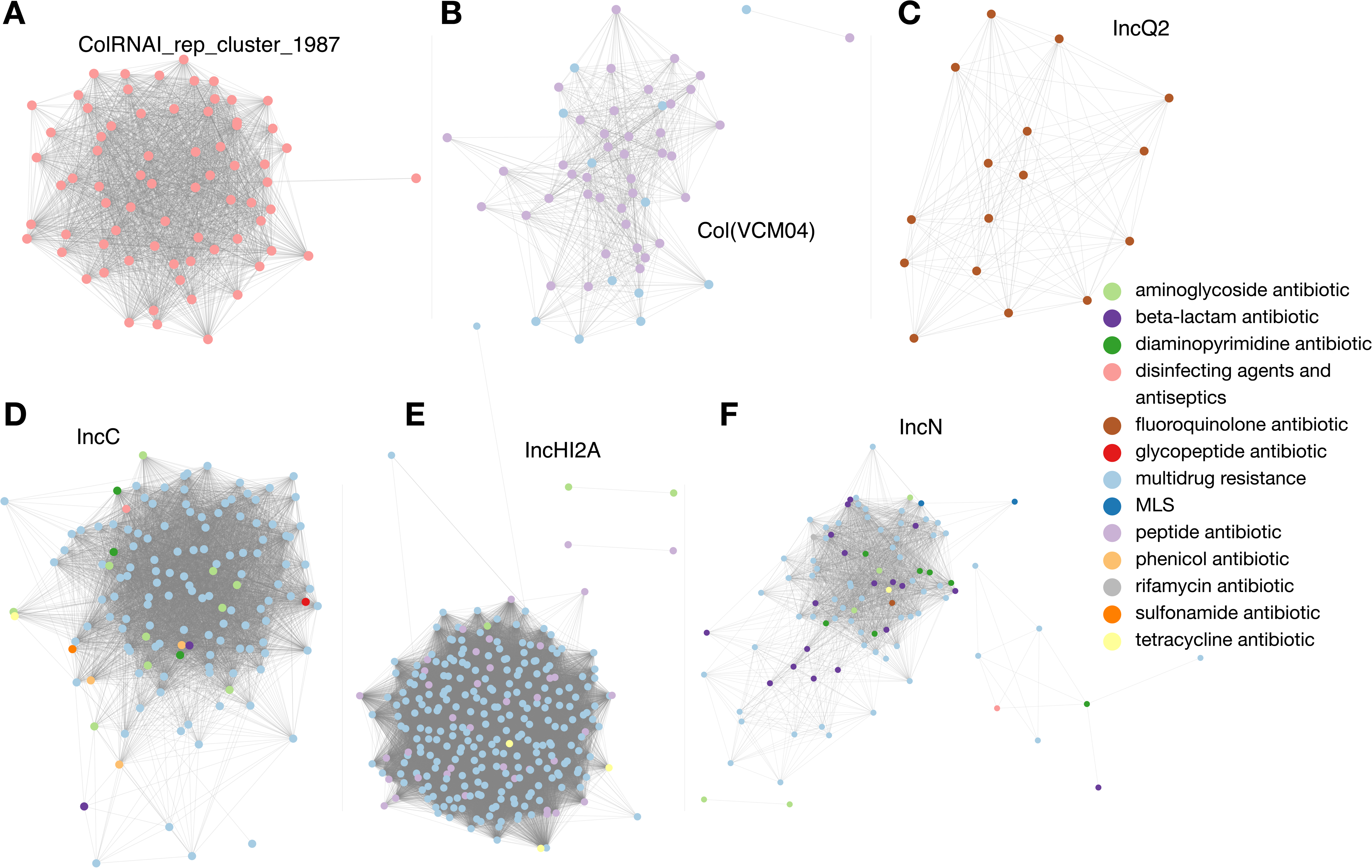
Genetic similarity networks of specific plasmid replicon types and their associated antibiotic resistance gene (ARG) profiles. Each node represents an individual plasmid, and edges connect plasmids with high genetic similarity. Nodes are colored according to the functional class of the carried antibiotic resistance genes (ARGs) as defined by the CARD database.

By mapping the ARGs to plasmids, we observed that specific plasmid types, such as ColRNAI_rep_cluster_1987, Col(VCM04), and IncQ2, exhibited strong associations with sulfonamide, peptide, and glycopeptide antibiotics, respectively (Figure 5A-C and Table 1). Specifically, the ARGs on ColRNAI_rep_cluster_1987 plasmids were disinfectant resistance genes *qacG*, while those on Col(VCM04) were primarily *mprF*. The IncQ2 plasmids were mainly associated with the *qnrS2* (Supplementary Table S8). In contrast, such distinct gene-replicon associations were rarely observed in plasmids harboring multidrug resistance, which often exhibited more complex and heterogeneous ARG compositions (Figure 5D-F, Table 1).

In terms of host distribution, the *qacG* genes associated with ColRNAI_rep_cluster_1987 were identified across 16 species, with 34 of the 71 total plasmids (47.9%) found in *Serratia nevei* (Supplementary Table S8). The *mprF* genes associated with Col(VCM04) spanned 14 species, with 40 of the 56 total plasmids (71.4%) originating from the *Citrobacter* genus. The *qnrS2* genes associated with IncQ2 were exclusively identified in *Leclercia adecarboxylata*.

## DISCUSSION

Across 6,895 Enterobacteriaceae genomes, we show that plasmid-borne antibiotic resistance genes (pARGs) and other plasmid genes exhibit similar species-level gain rates (Figure 1A). However, the expansion and reduction rates of ARGs, which depend on species, are much more diverse than those of other plasmids. Elevated expansion and reduction may reflect the activity of gene-level mobile elements such as IS26 translocatable units (23, 24) and integron cassette rearrangements (25, 26), as well as selective amplification events that duplicate individual genes without affecting entire plasmids (27). Besides, recent study revealed that 77% of studied ARG-carrying plasmids contain insertion sequences, facilitating inter-plasmid resistance gene transfer (7). And the replication-control mutations (e.g., *copA*) may also elevate plasmid copy number and increase ARG transfer rates (11)

Beyond species-specific variation in expansion and reduction, which likely reflects differing balances between antibiotic-driven selection and fitness costs (28, 29), we observed that bacterial lineages harboring cARGs acquired pARGs at 2.47-fold higher and 3.7-fold lower rates than their sister clades (Figure 6B, Supplementary Table S6). Rather than implying direct causation, the lack of temporal resolution in static genomic data precludes a definitive conclusion regarding the directionality of this evolutionary process. We consider two distinct evolutionary scenarios to explain this association. In one scenario, chromosomal integration acts as a prerequisite for further resistance accumulation. By fixing critical resistance genes into the core genome, lineages ensure vertical inheritance and minimize the metabolic burden associated with maintaining plasmids. This stabilized genetic background may facilitate the subsequent uptake of a broader plasmid repertoire. Alternatively, chromosomal integration may be the evolutionary consequence of intense plasmid flux under sustained antibiotic selection. High rates of pARG acquisition increase the intracellular density of resistance genes, elevating the statistical probability of stochastic transposition to the chromosome via mobile elements such as IS26 or integrons (24, 30–33). In this scenario, cARGs represent the consequence of a highly dynamic resistance mobilome. In any case, from a long-term evolutionary perspective, cARGs can serve as genomic markers for bacterial lineages prone to accumulate plasmid-mediated ARGs.

cARG-containing clades have a higher plasmid burden, with a median of 2.64 plasmids, compared to 1.66 in sister clades (Supplementary Table S7). Since the median sharing ratio of pARG-associated replicons with sister clades is 0%, these plasmids were likely acquired independently after the lineages diverged, rather than inherited from a common ancestor. This independent acquisition, combined with significantly lower rates of plasmid gene loss (Figure 3B), suggests that these clades are particularly effective at obtaining and retaining the environmental mobilome. The presence of more plasmids increases the physical proximity of different pARGs within the same cell. This co-localization creates the conditions necessary for recombination between different plasmid backbones or the assembly of new resistance islands (34). Consequently, these clades may be important sites for the exchange of ARGs. The newly formed elements are subsequently stabilized by reduced loss rates, ensuring persistent multidrug resistance and contributing to the dissemination of these traits to the broader bacterial population.

Finally, we manually selected few Count-inferred events (posterior probability >0.90) and verified whether the events matched previous reported outbreaks (Supplementary Table S9). We identified a clade of 24 genomes from France that acquired the *blaCTX-M* gene family at node N13 of the *Enterobacter quasihormaechei* species tree. These genomes were assigned to ST873 (using the ecloacae scheme), matching a previously documented *blaCTX-M-15* positive ST873 outbreak in French hospitals (35). Similarly, a gain event of the *mcr* phosphoethanolamine transferase family was inferred at node N22 of the *Escherichia fergusonii* species tree, involving 10 Chinese isolates. These genomes were assigned to the novel ST16672 type (using the ecoli_achtman_4 scheme), for which no known clinical outbreaks have been reported to date. However, by looking at their isolation locations, this gain event aligns with the landmark emergence of plasmid-mediated mcr-1 in Chinese animal and human populations (36). Furthermore, we identified multiple *sul* gain events across diverse species that align with regional antimicrobial resistance (AMR) trends. For instance, a gain at *Salmonella enterica* node N9 is consistent with the high *sul* prevalence (34.37%) reported in southern Brazil (37). Similarly, a gain at *Salmonella arizonae* node N34 matches documented MDR outbreaks in U.S. turkey operations (38), while a gain at *Edwardsiella anguillarum* node N5 reflects the high detection frequency of *sul* genes in fish pathogens along the Fujian coast of China (39).

### Limitations

This study focused exclusively on plasmid-associated genes, excluding other mobile elements such as Integrative and conjugative elements (ICEs) and integrative mobilizable elements (IMEs), which also mediate chromosomal integration and excision of ARGs (40–42). Bacteriophages further can contribute to ARG dissemination through transduction and phage–plasmid hybrids (43). Consequently, our observations represent only a subset of the overall resistance mobilome. Moreover, we lacked ecological information about antibiotic exposure histories of the analyzed organisms. Expanding future analyses to include additional mobile genetic elements and ecological datasets would provide a more comprehensive understanding of ARG dissemination in natural environments.

### Conclusion

Our findings highlight a duality in plasmid-mediated resistance evolution in Enterobacteriaceae. At the level of whole gene families, pARGs follow the same turnover dynamics as do other plasmid genes, indicating that resistance dissemination largely reflects plasmid-level processes rather than ARG-specific rules. At the level of copy-number changes, however, pARGs show elevated expansion and reduction with strong lineage dependence and minor drug-class effects. Such species-driven plasticity suggests recurrent expansion-reduction cycles as a flexible resistance strategy. Moreover, chromosomal resistance content serves as a genomic marker of a lineage’s potential for plasmid-mediated resistance accumulation on evolutionary timescales.

## METHODS

### Genome Collection and Gene Annotation

We parsed the Genome Taxonomy Database (GTDB; https://gtdb.ecogenomic.org) release 220 (44) to identify genomes belonging to Enterobacteriaceae. For species with more than 300 genomes, we randomly selected 300 representatives with random seed 19940421. We applied MIMAG criteria (45) with completeness >99%, contamination <1%, mean contig length >5,000 bp, and contig count <500 to filter high-quality genomes. To reduce computational cost, species with more than 100 genomes were subsampled to retain at most 100 genomes (random seed 19940421). Only species with more than 10 genomes were considered for further analysis. The genomes that passed these filters were downloaded from the NCBI FTP site (https://ftp.ncbi.nlm.nih.gov). For sequence typing, the MLST software (https://github.com/tseemann/mlst) was used to against PubMLST database (46).

Bacterial genes were predicted using pyrodigal-gv v0.3.2 integrated in geNomad. Plasmids and their encoded genes were identified using geNomad v1.11.0 with database v1.9 and default parameters (47), as well as the Platon v1.7 accuracy model (48). we defined a gene as plasmid-derived only if it is independently identified by both geNomad and Platon. Antibiotic resistance genes were identified with Resistance Gene Identifier (RGI) v6.0.5 using protein sequences with “include_nudge” option and default parameters, CARD version 4.0.1 (22). Pan-genomes of each species were constructed using the orthology-based clustering method OrthoFinder v2.5.5 (49) following established guidelines (50).

After applying these criteria, 6,895 complete or nearly complete genomes belonging to 159 Enterobacteriaceae species (sensu GTDB) were included in the analysis (Supplementary Table S1), of which 80 contained antibiotic resistant genes on plasmids (Supplementary Data).

### Applying Phylogenetic Birth-Death Model for Gene Gain and Loss Inference

From the orthologous gene families and species trees generated by OrthoFinder, we extracted plasmid orthologous gene families. An orthologous gene family was considered plasmid-associated if the proportion of plasmid genes within that orthologous gene family exceeded 70% of all genes in the family. Species trees were pruned to retain only genomes contributing to the filtered orthologous gene families.

Gene gain and loss dynamics were quantified using the phylogenetic birth-death model implemented in Count v10.04 (21). This probabilistic framework models the evolution of gene family sizes along phylogenetic trees as a continuous-time Markov process. The model accounts for three fundamental processes: gene gain (κ), gene loss (μ), and gene duplication (λ). A gene family of size n changes according to the following rates: the family decreases at rate nμ due to individual gene losses, increases through duplication at rate nλ, and gains new genes at rate κ through horizontal transfer or other acquisition mechanisms.

Note that the biological processes underlying gene gain, loss, expansion, and reduction can be interpreted as follows: 1) Gain (0 to 1): A species acquires a new plasmid carrying a previously absent resistance gene family, or a new gene family integrates into an existing replicon via horizontal gene transfer. 2) Loss (n to 0 for n > 1): The total loss of a plasmid harboring all copies of a gene family, or the complete deletion of a gene family from all replicons within the cell. 3) Expansion (n to m for m > 1): An increase in intra-replicon copy number via tandem duplication, which is often mediated by insertion sequences; the acquisition of additional homologous copies from external sources into existing plasmids; an increase in plasmid copy number via replication-control mutations; or the translocation and subsequent duplication of pARG onto the chromosome or other co-resident plasmids. 4) Reduction (n to m for n > m ≥ 1): Loss of some duplicated copies of a gene family while retaining at least one copy, or loss of one plasmid in a cell harboring multiple replicons with the same gene family. These scenarios illustrate the most probable biological drivers captured by the model. However, they are not an exhaustive representation of all complex scenarios that may occur in Enterobacteriaceae genomes.

For each species dataset, we performed iterative maximum likelihood optimization to estimate birth-death model parameters using whole bacterial genomes (51). We implemented a systematic search strategy starting with initial parameter values of κ=μ=λ=1 and incrementally exploring the parameter space up to κ, μ, λ ≤ 4. This resulted in a total of 10 optimization rounds. The iterative approach followed the pattern: if all parameters are equal, increase loss rate (μ); if duplication and gain rates are equal, increase duplication rate (λ); otherwise increase gain rate (κ). Each round uses the previous round’s results as starting values. The optimization utilized maximum likelihood estimation with maximum paralogs of 200, optimization tolerance of 0.1, and minimum lineages of 2.

After parameter optimization, we calculated posterior probabilities for plasmid-borne ancestral gene family states and evolutionary events using the Posteriors module of Count. This reconstructs the evolutionary history of each gene family, providing probabilistic estimates for gain, loss, expansion, and reduction events along each branch. Count analysis was performed separately for each species dataset.

### Gene Gain and Loss Data Integration and Rate Calculation

An orthologous gene family was defined as an ARG family if at least 50% of its member genes were identified as ARGs by RGI. Branch lengths were extracted from the original species trees and combined with corresponding Count posterior probabilities. For rate calculations, we summed gain, loss, expansion, and reduction events across all branches within each orthologous gene family and divided by the corresponding total branch length. Only branches with branch lengths > 1e-12 were included to avoid computational artifacts from extremely short branches. Evolutionary rates were calculated per species per orthologous gene family as:

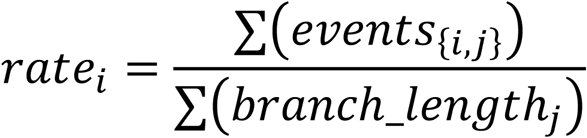

where *events*_{*i,j*}_ is the number of events for orthologous gene family *i* on branch *j*, and *branch*_*length*_{*j*}_ is the length of branch *j* in substitutions per site.

### Comparative Analysis of Evolutionary Rates

For comparative analysis between ARG and non-ARG plasmid gene families, data were aggregated by species, event type, and ARG status. Median rates were calculated within each group, and the data were restructured into a paired format where each species served as a matched pair with corresponding median rates for ARG and non-ARG families. Paired Wilcoxon signed-rank tests compared ARG versus non-ARG families within species to control for species-specific evolutionary factors. Additionally, gain/loss and expansion/reduction ratios were calculated per species to assess the relative balance of these complementary processes. Multiple testing correction was performed using the Benjamini-Hochberg procedure. Effect sizes were quantified as median log2-fold changes, and statistical significance was assessed at p < 0.05 and q < 0.05 (FDR-adjusted p).

To examine rate variation across antibiotic resistance classes, drug class annotations were extracted from RGI output. For ARG families associated with multiple drug classes, evolutionary rates were assigned to each corresponding class. Evolutionary rates were then visualized across different species and drug class within Enterobacteriaceae. The concordance between event types across species was assessed by compared median per-species rates using Spearman’s rank correlation with permutation tests (5,000 iterations, random seed = 19940421) to obtain empirical p-values.

To evaluate the relative contributions of species background and drug class to pARG turnover, variance partitioning was performed using Ordinary Least Squares (OLS) models. Evolutionary rates (*y*) were log-transformed as ln(*y* + 10^−12^) to satisfy normality assumptions. We compared the explanatory power of two categorical models, Species-driven model, *ln*(*y* + *∈*) ∼*β*_0_ + *β_i_* · *Species_i_* + *∈*, and a drug-class-driven model, *ln*(*y* + *∈*) ∼*β*_0_ + *β_i_* · *Drug*_*class_j_* + *∈*, where Species and drug class represent categorical predictors for bacterial lineages and antibiotic classes, respectively. The coefficient of determination (R^2^) was used to quantify the proportion of total variance explained by each factor.

### Comparison of Plasmid Gene Dynamics Between ARG-containing and Sister Clades

To assess whether the presence of antibiotic resistance genes in bacterial chromosomes influences the evolutionary dynamics of plasmid-borne genes, we identified pairs of phylogenetic clades that differed in the chromosomal presence of an ARG family. For each combination of species tree and ARG family, we scanned for internal nodes where ≥80% of descendant genomes harbored the specific ARG family in their chromosome (cARG-containing clade), while <20% of genomes in the corresponding sister clade contained that ARG family. Only clade pairs with at least two genomes in each clade were retained for analysis. A representative case involving Citrobacter portucalensis is illustrated in Supplementary Figure S8.

For each identified clade pair, we calculated the gain and loss rates of plasmid genes, distinguishing between plasmid-borne ARGs and other plasmid genes. For each orthologous gene family, rates were computed as:

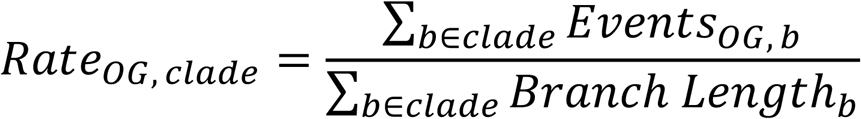

where events were summed across all branches within each clade, normalized by the total branch length of that entire clade. Plasmid ARGs that matched the chromosomal ARG family used to define the clade were excluded from the analysis. The median rate across all orthologous gene families within each gene class (ARGs or non-ARGs) was then used as the representative value for each clade.

Event rates were analyzed using linear mixed-effects models on the log scale, with gain and loss modeled separately; the response was log(*rate* + ε), where ε = 0.1 × min(non-zero rate) within the corresponding event type. Fixed effects were clade type (ARG-containing vs sister clade), gene class (plasmid ARGs vs other plasmid genes), and their interaction; random effects were species, resistance gene family, and the clade pair linking each ARG-containing clade with its sister (implemented as a random intercept on Clade ID). Models were fitted using restricted maximum likelihood and restricted to paired observations with both clade types available within a given gene class. Post hoc contrasts from estimated marginal means assessed (i) ARG-containing vs sister differences within each gene class and (ii) the clade-type × gene-class interaction. In addition to the overall model, we performed family-specific analyses. For each resistance gene family with ≥ 4 available paired clades, we re-fit the model within that family using the same fixed-effects structure and random intercepts for species and clade pair, but omitting the family random effect (not identifiable within a single family). Estimates and 95% CIs were back-transformed to the original rate scale, with two-sided tests and Benjamini–Hochberg adjustment for multiple contrasts. Analyses were performed in R v4.4.0 using lme4 v1.1.35.5 (52), lmerTest v3.1.3 (53), and emmeans v 1.11.2.8.

To evaluate the robustness of our sister-clade analysis, we performed a sensitivity analysis by testing a gradient of clade-defining thresholds for cARG frequency: 70%/30%, 80%/20% (the primary threshold used in this study), 85%/15%, and 90%/10%. These alternative thresholds were applied to verify whether the observed associations between cARG presence and pARG gain/loss rates were sensitive to the stringency of clade selection. The fold-change in pARG gain rates and the decrease in pARG loss rates were assessed across all thresholds using the linear mixed-effects model structure described above.

The usage levels of ARG-associated antibiotics in clinical (human, Spain, 2023) and agricultural (veterinary, Spain, 2022) data were obtained from the One Health Trust’s ResistanceMap (https://resistancemap.onehealthtrust.org/) and the European Medicines Agency’s (EMA) ESVAC report (https://www.ema.europa.eu/), respectively, and were standardized into the CARD drug class.

### Plasmid Unit Reconstruction and Similarity Network Analysis

To accurately determine plasmid numbers and mitigate overestimation, mob_recon (MOB-suite v3.1.9) (54) was employed to reconstruct plasmids across the entire dataset. In this section, each reconstructed plasmid was treated as an individual genomic unit. Only plasmid groups (primary clusters) comprising at least ten independent units and a consistent replicon type were included in the analysis. Genomic similarity within each group was assessed by calculating Mash distances, excluding edges between plasmid pairs with distances exceeding 0.025. The resulting similarity networks were visualized using Cytoscape v3.10.2 (55).

Resistance genes conferring resistance to monobactams, carbapenems, cephalosporins, or penicillins were grouped into a single beta-lactam category for the drug-class analysis. Similarly, the MLS group was used to encompass macrolides, lincosamides, pleuromutilins, and streptogramins (A and B).

## ACKNOWLEDGEMENTS

Yue Liu is supported by the State Key Laboratory for Diagnosis and Treatment of Infectious Diseases (No. 202506) and the ZJU100 Young Professor Start-up Fund from Zhejiang University of China. Yang Liu is supported by the China Postdoctoral Science Foundation (No. 2025M783825).

## DECLARATION OF GENERATIVE AI AND AI-ASSISTED TECHNOLOGIES IN THE WRITING PROCESS

During the preparation of this work, the author(s) used ChatGPT and Gemini to polish the language. After using this tool, the authors reviewed and edited the content as needed and took full responsibility for the content of the publication.

## ETHICS APPROVAL

There are no ethical concerns in this study.

## DATA AVAILABILITY

Supplementary material is available for this article. Also, the supplement data, and core script are available on Figshare under the DOI: 10.6084/m9.figshare.30186307.

## AUTHOR CONTRIBUTIONS

**Yang Liu:** conceptualization, bioinformatics analysis, writing – original draft, writing – review & editing. **Yue Liu:** writing – review & editing, funding acquisition.

