## Supplemental Figure for "Gain and loss of plasmid-borne antibiotic resistance genes are associated with chromosomal resistance presence in Enterobacteriaceae"

### Supplementary Figure Legends

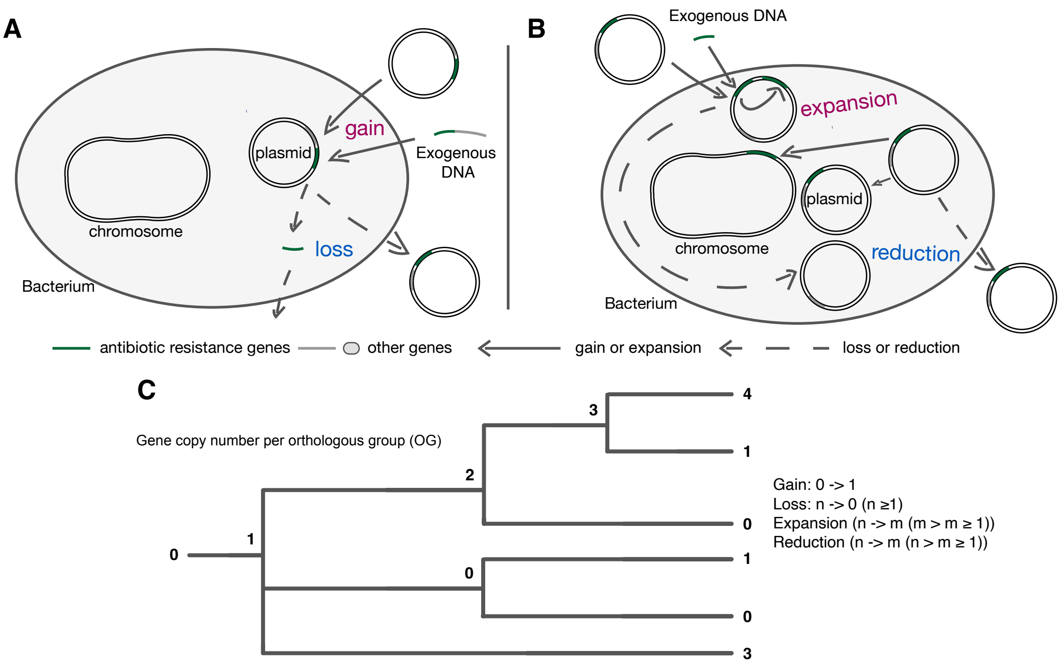


**Figure S1. Definitions of the four plasmid gene evolutionary processes.** (A) gene gain and loss. Gain is defined as the acquisition of a new orthologous group (OG); Loss is defined as the complete disappearance of an existing OG (n to 0). (B) expansion and reduction. Expansion is defined as an increase in gene copies within an existing OG (n to m, m > n ≥ 1); Reduction is defined as a decrease in gene copies while maintaining at least one copy (n to m, n > m ≥ 1). (C) Numerical definitions. Schematic of copy-number state transitions per OG for each process.

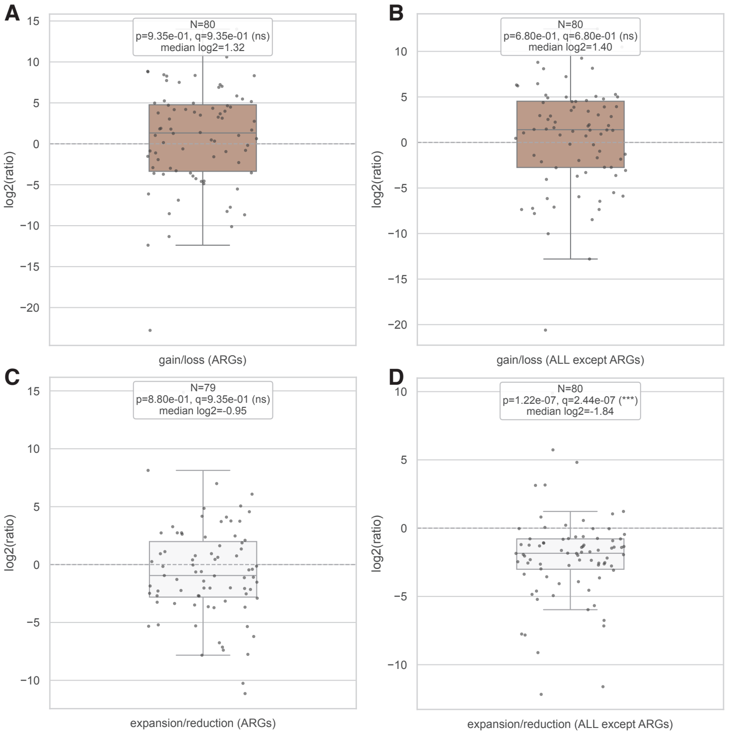


**Figure S2. Ratio of gene copy number across species.** (A) $Log_2$ ratio of gain to loss rates for ARGs. (B) $Log_2$ ratio of gain to loss rates for other plasmid genes. (C) $Log_2$ ratio of expansion to reduction rates for ARGs. (D) $Log_2$ ratio of expansion to reduction rates for other plasmid genes. Each point represents one Enterobacteriaceae species. Box plots show median and quartiles. Gene families were defined as orthologous groups (OGs).

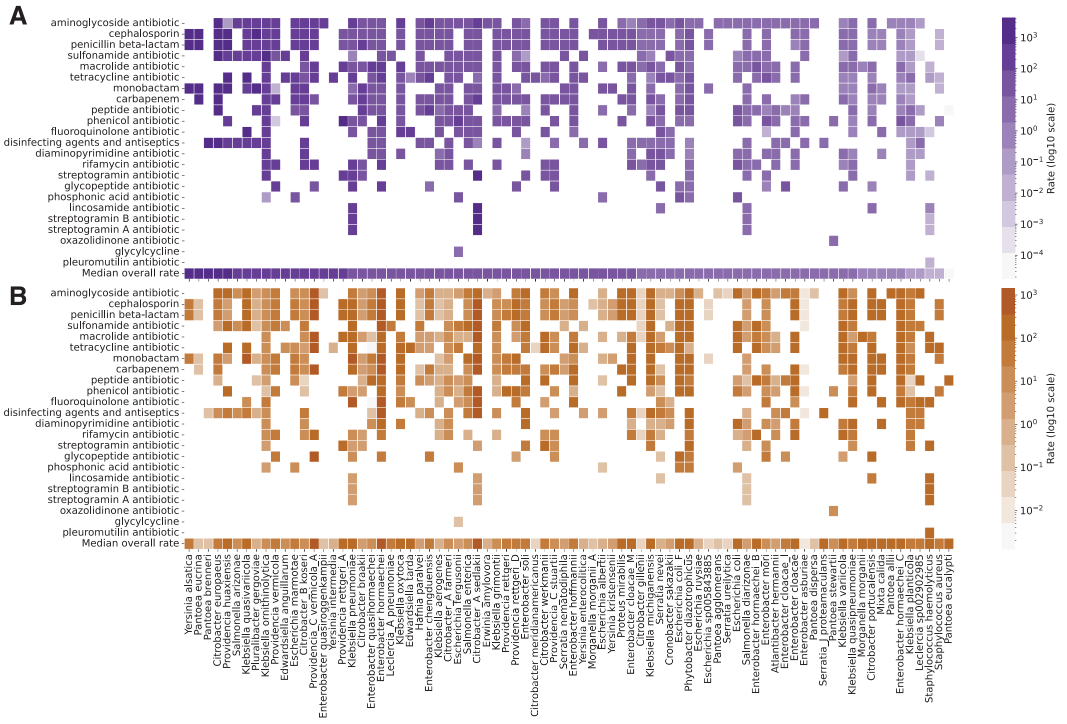


**Figure S3. Gain and Loss rate patterns across antibiotic resistance classes and Enterobacteriaceae species.** Heatmaps showing evolutionary rates for different antibiotic resistance classes across bacterial species. (A) Gain rates. (B) Loss rates. Rows represent Enterobacteriaceae species, and columns represent CARD-defined antibiotic resistance classes. Color intensity indicates log10 scaled rate magnitude, with darker colors representing higher rates. White cells indicate absence of the ARG class in the species.

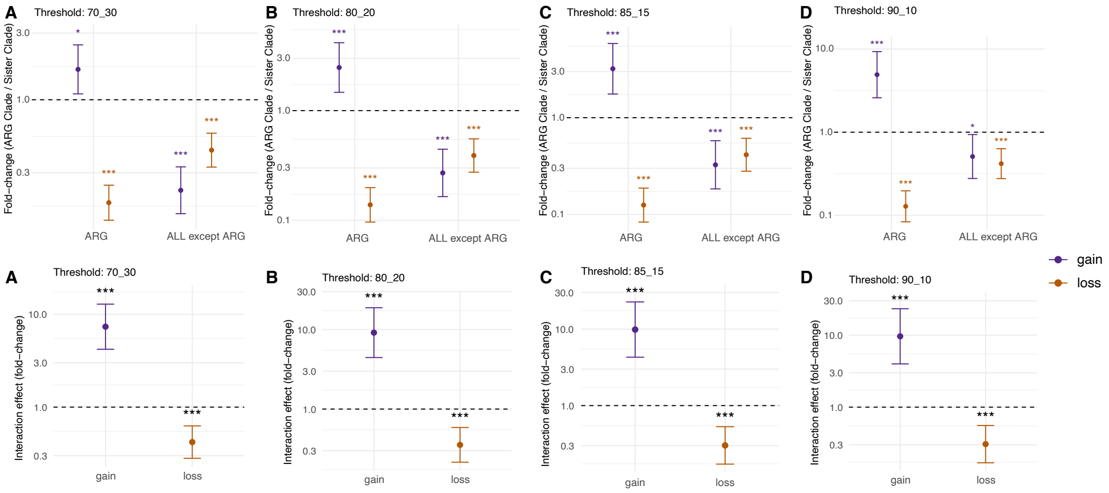


**Figure S4. Robustness of pARG gain and loss dynamics across different sister-clade thresholds.** Sensitivity analysis using various thresholds for defining ARG-containing and sister clades. Thresholds (cARG/sister) are specified for each panel: (A) 70/30, (B) 80/20, (C) 85/15, and (D) 90/10. Fold-change comparison (ARG clade / Sister clade) for gain (purple) and loss (orange) rates of plasmid ARGs versus other plasmid genes (ALL except ARG). The dashed line at y = 1 indicates no difference between clades. Statistical significance indicators: ***p < 0.001. Error bars represent 95% confidence intervals.

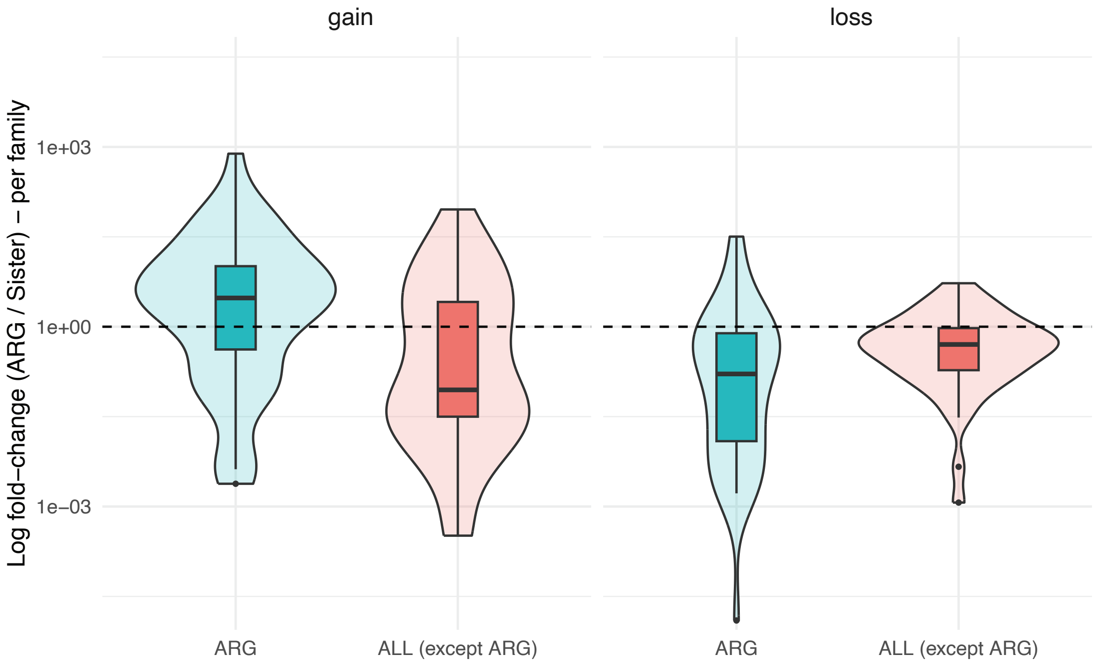


**Figure S5. Comparison of evolutionary rates between ARG-containing clades and their sister clades in per-family analyses.** Y-axis shows the fold-change ratio (ARG clade / Sister clade) of plasmid ARGs versus other plasmid genes. Dashed line at y=1 indicates no difference between clades. Violin plots show data distribution, and box plots indicate the median and interquartile range.

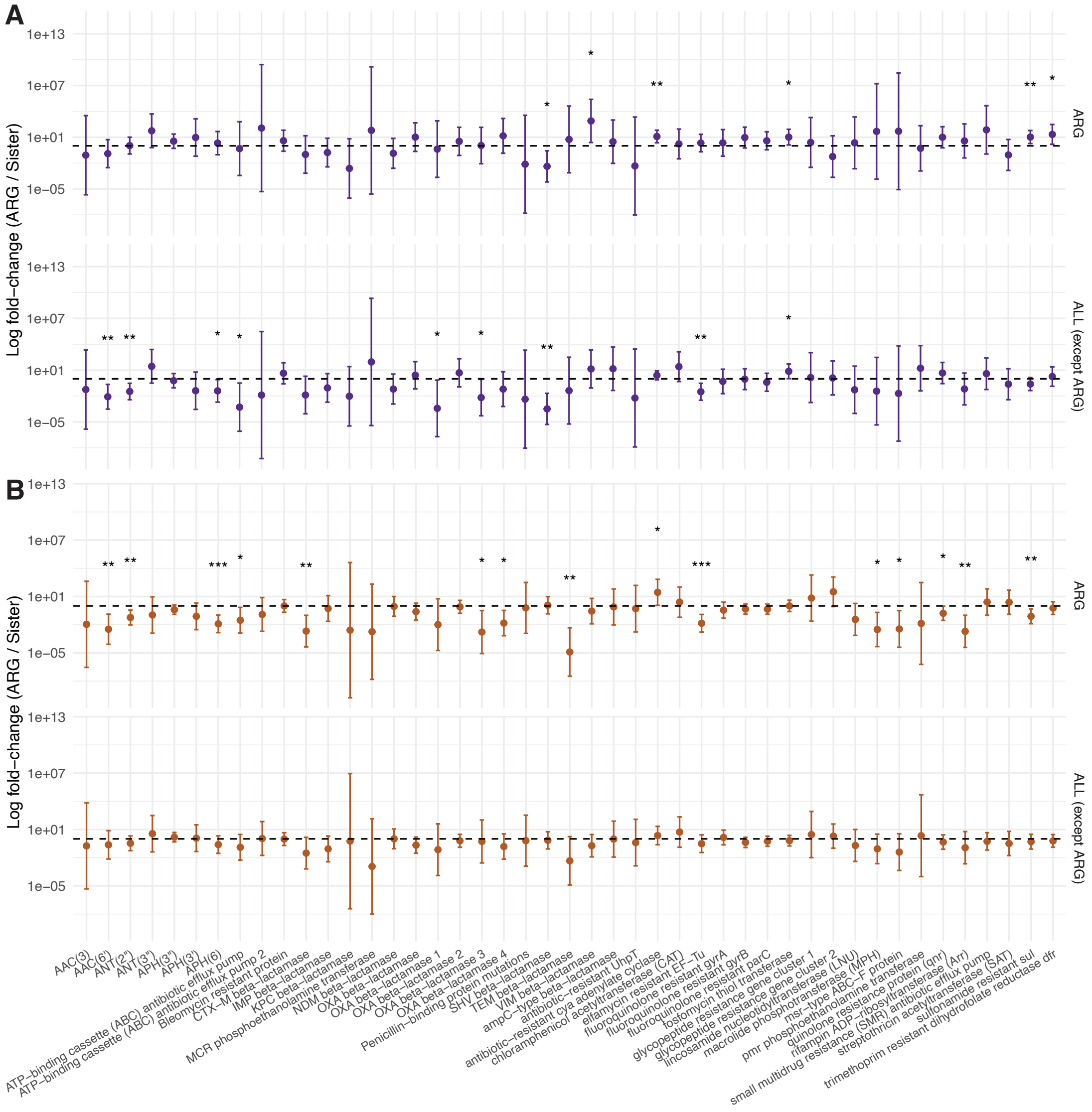


**Figure S6. Per-family comparison of evolutionary rates between ARG-containing clades and their sister clades.** (A) Gain rates and (B) loss rates are shown for individual antibiotic resistance gene (ARG) families. A dashed line at y = 1 indicates no difference between clades. Asterisks denote statistical significance (* $p < 0.05$; ** $p < 0.01$; *** $p < 0.001$).

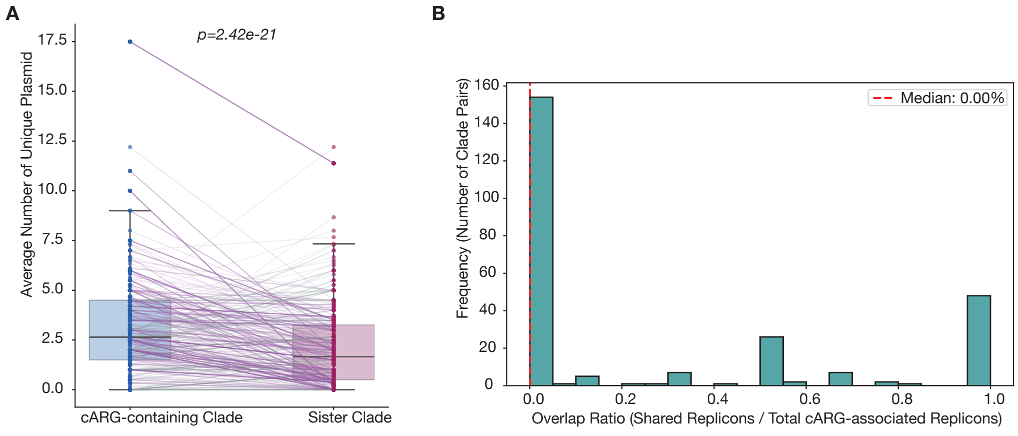


**Figure S7. Comparison of plasmid burden and replicon sharing between cARG-containing and sister clades.** (A) Pairwise comparison of plasmid burden, Wilcoxon test. (B) The histogram shows the distribution of shared plasmid replicons between cARG clades and their sister clades.

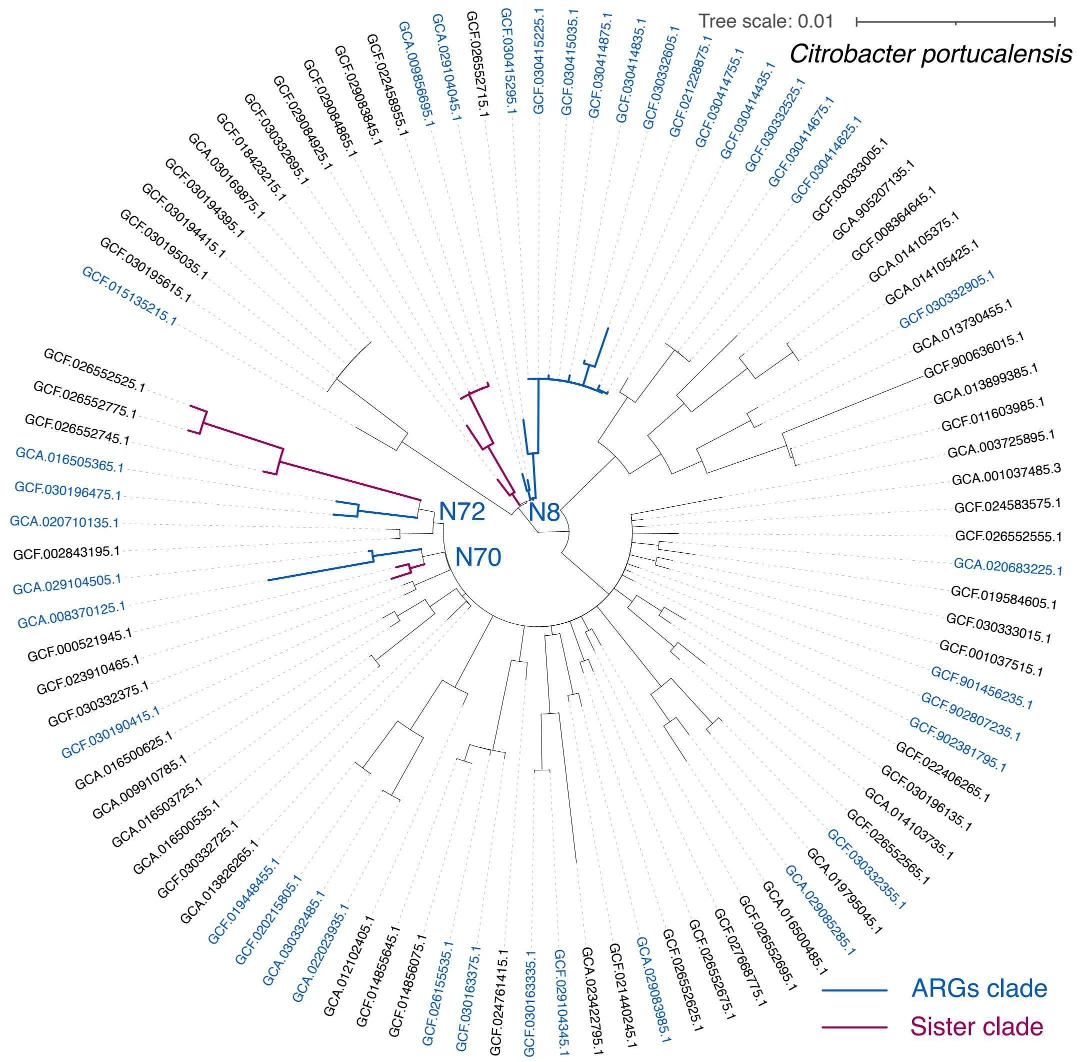


**Figure S8. An example of a phylogenetic tree with annotated ARG-containing and sister clades.** Gene family: fluoroquinolone resistant *gyrA*. Species representative genome accession: GCF_002042885.1. Blue text indicates that there is at least one corresponding ARG in the genome.
